# Predicting stable binding modes from simulated dimers of the D76N mutant of *β*2-microglobulin

**DOI:** 10.1101/2021.07.14.452361

**Authors:** Nuno F. B. Oliveira, Filipe E. P. Rodrigues, João N. M. Vitorino, Rui J. S. Loureiro, Patrícia F. N. Faísca, Miguel Machuqueiro

## Abstract

The D76N mutant of the *β*_2_m protein is a biologically motivated model system to study protein aggregation. There is strong experimental evidence, supported by molecular simulations, that D76N populates a highly dynamic conformation (which we originally named I_2_) that exposes aggregation-prone patches as a result of the detachment of the two terminal regions. Here, we use Molecular Dynamics simulations to study the stability of an ensemble of dimers of I_2_ generated via protein-protein docking. MM-PBSA calculations indicate that within the ensemble of investigated dimers the major contribution to interface stabilization at physiological pH comes from hydrophobic interactions between apolar residues. Our structural analysis also reveals that the interfacial region associated with the most stable binding modes are particularly rich in residues pertaining to both the N- and C-terminus, as well residues from the BC- and DE-loops. On the other hand, the less stable interfaces are stabilized by intermolecular interactions involving residues from the CD- and EF-loops. By focusing on the most stable binding modes, we used a simple geometric rule to propagate the corresponding dimer interfaces. We found that, in the absence of any kind of structural rearrangement occurring at an early stage of the oligomerization pathway, some interfaces drive a self-limited growth process, while others can be propagated indefinitely allowing the formation of long, polymerized chains. In particular, the interfacial region of the most stable binding mode reported here falls in the class of self-limited growth.

**Graphical Abstract:** 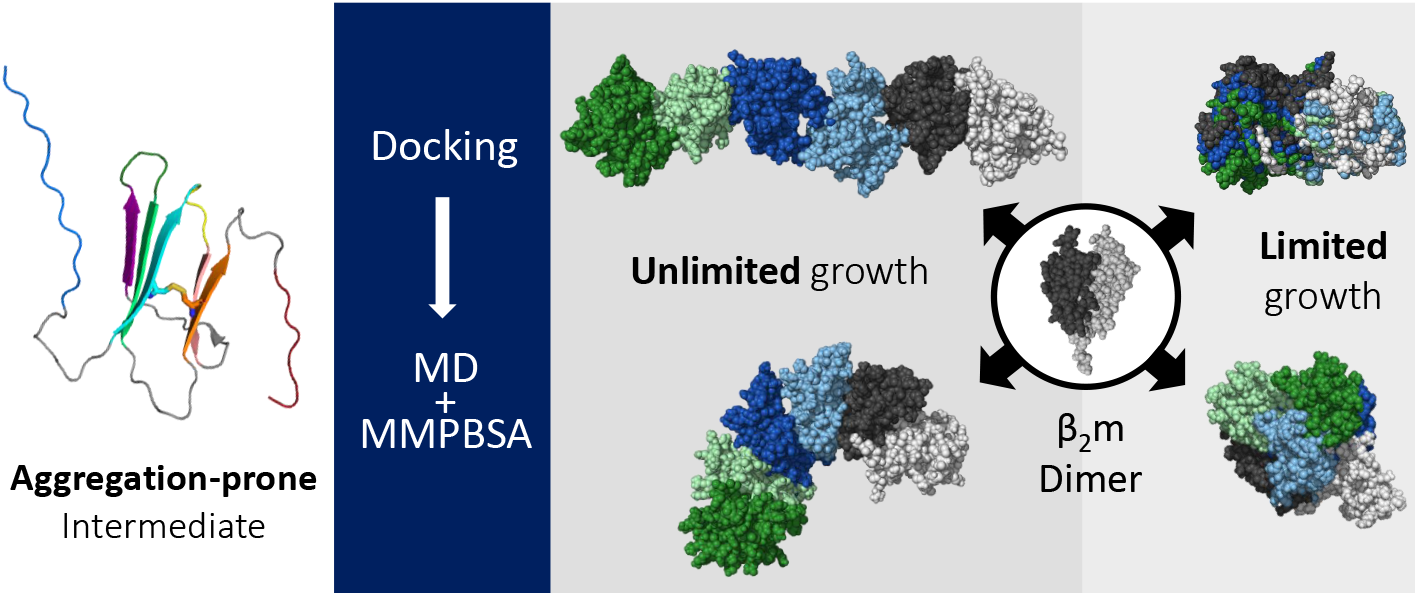

**Highlights:**

- The D76N mutant of protein *β*_2_m populates an aggregation-prone monomer (I_2_) with unstructured terminal regions
- Molecular Dynamics simulations and MM-PBSA calculations indicate that dimers of I_2_ are stabilized by hydrophobic interactions
- The N- and C-terminal regions, together with the BC- and DE-loops are prevalent in the most stable dimer interfaces, while the CD- and EF-loop appear in the less stable ones
- The most stable dimer interface has a limited potential to oligomerize in the absence of structural rearrangement

## 1. Introduction

Beta-2 microglobulin (β_2_m) is a small protein whose native structure exhibits a typical immunoglobulin fold. The 99 residues that compose the wild-type (*wt*) form are arranged into two sheets of anti parallel beta-strands forming a sandwich-like structure. One of the sheets comprises strands A-B-E-D, while the other is composed by strands C-F-G. The native structure is stabilized by a disulfide bridge linking residues Cys25 (strand B) and Cys80 (strand F) [1, 2] (Figure 1A-B).

**Figure 1:**
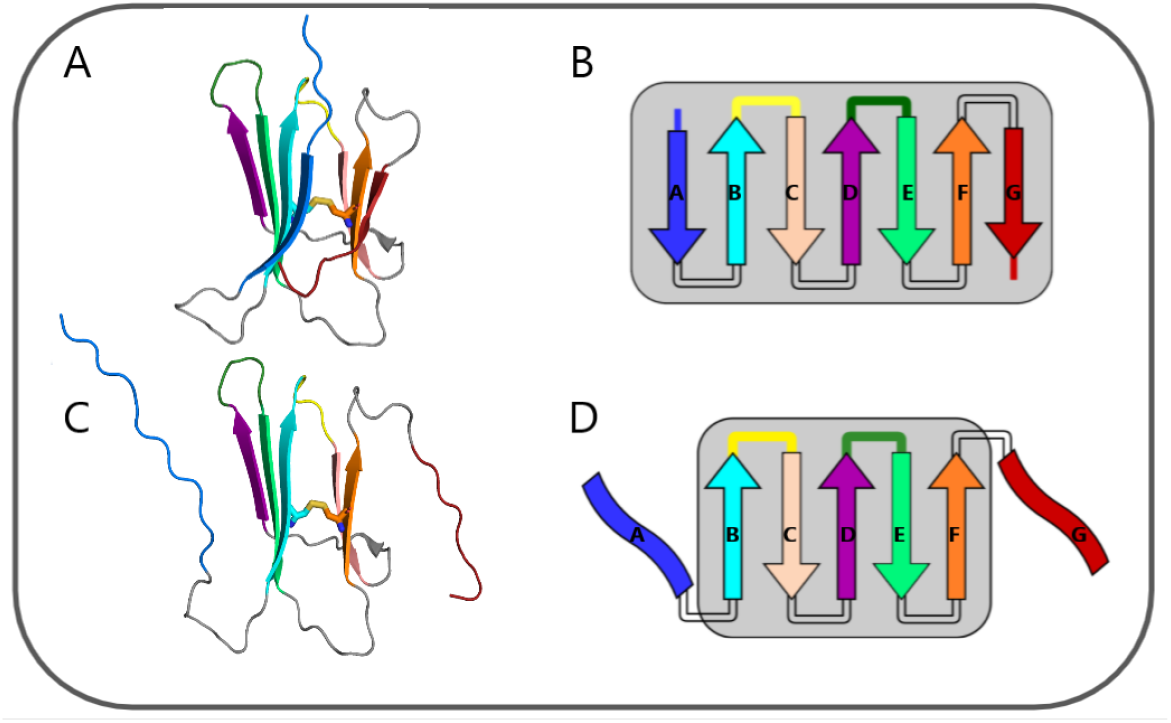
Three-dimensional cartoon representation of the native structure of D76N (pdb id: 2YXF[2])(A) and the schematic representation of its secondary structure (B). The I_2_ intermediate state populated by D76N, which features a well-preserved core and two unstructured and decoupled termini (C, D). The N- and C-termini are coloured in blue and red, respectively. The BC- and DE-loop are colored yellow and green, respectively.

The *wt* form is the causing agent of dialysis related amyloidosis (DRA), a conformational disorder associated with the formation of amyloid fibrils in the osteoarticular system of individuals undergoing long-term (>10 yrs) hemodialysis [3]. The *ex-vivo* amyloid fibrils of patients with DRA are composed of the wt form, and, to a lesser extend (30%), of a truncated structural variant lacking the six N-terminal residues, which is termed ΔN6 [4]. The wt form does not aggregate *in vitro* under physiological conditions, while the ΔN6 does [5, 6]. For this reason, it has been extensively used as a model system to study *β*_2_m aggregation. However, the biological significance of ΔN6 lacks consensus because it is not clear if the proteolytic cleavage of the terminal hexapeptide occurs before, or after, fibril assembly [7].

More recently, the D76N mutant, was identified as being the culprit of a rare systemic amyloidosis pathology affecting visceral organs (liver, kidneys etc.) [8], which readily aggregates *in vitro* under physiological conditions [9]. This single point mutant thus appears to be a suitable model system to study the details of *β*_2_m aggregation mechanism.

The small size of *β*_2_m, together with its biomedical significance, renders it an ideal target for *in silico* investigations framed on molecular simulations, and a plethora of interesting results has been reported in the literature (reviewed in [10]). In particular, some of us developed a three-stage approach that integrates information obtained from complementary simulation methodologies [11, 12, 13], to provide a probabilistic, structurally-resolved picture of the initial phase of the *β*_2_m aggregation mechanism, which starts with the formation of dimers [14]. Specifically, we use discrete Molecular Dynamics of a full atomistic structure-based model [15, 16] to explore the folding space of the monomer and detect intermediate states with aggregation potential. The identified intermediate states are then used as starting conformations in constant-pH molecular dynamics simulations [17, 18, 19] to create ensembles of monomers representative of different environmental conditions. In the last stage of our procedure, we deploy an in-house protein-protein Monte Carlo ensemble docking method (MC-ED), to create ensembles of dimers whose statistical analysis allows predicting which regions of the monomer are more likely to self-associate into dimers, and identifying the so-called aggregation hot-spots, i.e., the residues that establish more intermolecular interactions in the interfacial region, being therefore critical to initiate the aggregation pathway.

In the particular case of the D76N mutant, our studies predict the existence of an aggregation-prone intermediate state, termed I_2_, that is structurally characterized by having two unstructured terminal regions, which are decoupled from (a well-preserved) protein core [12] (Figure 1C-D). We found that this intermediate is more likely to self-associate via the DE-loop and EF-loop under physiological conditions, with the termini becoming more relevant as adhesion zones under acidic pH [13]. The detachment and unfolding of the terminal regions predicted by our structure-based model is consistent with the fact that the D76N substitution breaks an important network of electrostatic interactions distributed over a large number of residues including both termini. This is in a rather good agreement with results based on ssNMR, which shows that D76N sparsely populates a highly dynamic conformation that exposes aggregation-prone regions as a result of a loss of *β*-structure at the terminal strands, i.e., an intermediate state that is topologically equivalent to I_2_ [20]. It has been suggested that the shear forces present in the extracellular fluid under physiological conditions may be enough to further unfold the intermediate termini and induce amyloid formation *in vivo* [21].

Our statistical analysis of the protein-protein interfaces based on MC-ED uses a full atomistic, but, nevertheless, rigid representation of the protein. This means, in particular, that it does not take into account the conformational changes that may occur upon binding, and which can go from local rearrangements of the side-chains, to large domain motions. Additionally, it uses a cost (or scoring) function based on square-well potentials that despite considering the three driving forces of protein self-association (shape, hydrophobic, and electrostatic complementarity that includes hydrogen bonds), does not explicitly take into account desolvation effects. Therefore, it provides a qualitative assessment on the stability of the interfaces of the dimers, i.e., of their binding energies. More precise methods, combining the atomic resolution of Molecular Mechanics with an implicit continuum solvent model, have been developed to quantitatively predict the binding energies of protein–protein complexes. In contrast to the docking scoring of single structures, an ensemble of complex conformations generated using Molecular Dynamics is evaluated in MM Poisson-Boltzmann/surface area (MM-PBSA) or MM generalized Born/surface area (MM-GBSA) approaches [22]. Both these methods have been shown to successfully predict protein-protein binding constants [23] and are particularly useful when the relative energy values between conformers are targeted since, in these cases, the elusive entropic contribution to the binding free energy mostly cancels out [24].

In the present work, we study in detail an ensemble of dimers obtained from monomers of the I_2_ intermediate state using the MC-ED method. In order to do so, we designed a protocol based on molecular dynamics simulations that relaxes the dimer conformation/configuration, and is followed by an evaluation of the dimer binding energy using a novel, in-house implementation of the MM-PBSA method. Having an ensemble of predicted dimer conformations, we sought to identify binding interfaces or binding modes (BMs) that conjugate energetic stability with unlimited growth, i.e, the ability to form long, elongated polymeric chains without having to undergo large structural rearrangements, and BMs that in the absence of large structural rearrangement are limited to grow into small oligomers. The results reported in the present work may have implications regarding the understanding of the aggregation mechanism of *β*_2_m based on the D76N mutant.

## 2. Methods

### 2.1. Molecular Dynamics simulations

We started by considering an ensemble of 221 dimers formed by monomers of the intermediate state I_2_, populated by the D76N mutant under physiological conditions (pH 7.2), which was obtained through the MCED method [13]. The molecular dynamics (MD) simulations of the dimers were performed using GROMACS 2018.6 [25], the GROMOS 54A7 force field [26, 27], and the SPC water model [28]. The full system used in the simulations comprises a dimer of two I_2_ monomers, ~20.000 water molecules, and 2 Na^+^ ions to keep the overall charge neutral. All ionizable residues in the protein were kept in their most probable protonation state at neutral pH[29], including histidines which were all kept neutral. In all production runs, the Particle-Mesh Ewald (PME) electrostatics [30, 31] was applied with a verlet scheme cutoff of 1.4 nm, a Fourier grid spacing of 0.12 nm, and an interpolation order of 4 (cubic). Van der Waals interactions were treated with a Lennard-Jones potential and were truncated above 1.4 nm. All bonds were constrained using the LINCS algorithm [32] for the protein while SETTLE was used for the water molecules [33]. The equations of motion were integrated every 2 fs with the neighbor lists being updated every 10 steps.

The dimer conformations were subjected to a classical protocol of energy minimization and MD initialization to ensure GROMOS 54A7 compatibility. A three-step protocol of energy minimization was performed consisting of 10.000 steps with the steepest-descent algorithm followed by 2.000 steps with the low-memory Broyden-Fletcher–Goldfarb–Shanno (l-BFGS) algorithm without constraints. Finally, the LINCS algorithm is activated on all bonds in a shorter third step (~100 steps) using again the steepest–descent algorithm. Any existent atomic clashes were removed in these three energy minimization steps. However, to start the temperature (NVT) and pressure (NPT) couplings in the MD initialization procedure, special attention to the position restraints is required and a three-step initialization protocol was also devised. The first step consisted on a 100 ps run in which the system temperature was raised to 310 K using the v-rescale thermostat [34] and a temperature coupling constant of 0.1 ps. Position restraints were applied to the C*α* atoms with a force of 1000 kJ mol^−1^ nm^−2^. In the second step (200 ps), we turned on the Parrinello-Rahman barostat [35, 36] to obtain an isotropic pressure of 1 bar with a pressure coupling constant of 0.5 ps and compressibility of 4.5 × 10^−5^ bar^−1^. In this step, the position restraints force constant was reduced to 100 kJ mol^−1^ nm^−2^. The third and final initialization step (200 ps) was focused on attenuating the position restraints, which now use a force constant of 10 kJ mol^−1^ nm^−2^. After this protocol, we performed a visual inspection step to identify conformations in which the MCED artificially introduced structural entanglements between monomers (Figure S1 of Supporting Information). Such structures were found in 9 dimer systems (1, 19, 73, 102, 104, 108, 161, 180 and 194), which were excluded from the present study. The remaining 212 dimer conformations were subsequently relaxed with unrestrained MD for 100 ns. In most cases the equilibrium properties converged relatively fast, but in a few cases the dimeric interfaces required longer equilibration times. Therefore, we performed equilibrium analyses using only the last 20 ns of each MD segment.

### 2.2. Structural analysis

All structural analyses of the dimers were carried out using the GROMACS package and other in-house tools. For the structural alignments and Root Mean Square Deviation (RMSD) calculations, only the the C*α* atoms from the protein core (namely, 23-27, 36-39. 51-55, 62-66. 78-82 of each monomer) were considered. In doing so, we discard spurious structural variability due to loops, and unstructured N- and C-terminal regions. The interfacial areas of the dimers were calculated using the SAS values of both monomers in the presence and absence of the partner [37]. This procedure was also applied to each residue individually, with the final area being normalized (converted into a percentage) by the maximum SAS area of each residue type observed in all simulations. Residues with interfacial area higher than 10%, were considered to be located at the interface. All error bars indicate the standard error of the mean. Images were rendered using PyMOL [38].

### 2.3. Structural clustering

Due to the large number of dimer configurations explored in this work, we applied a clustering protocol comprising 3 steps to reduce structural variability and try to ensure that unique dimers, representative of distinct binding modes, are captured and characterized. In the first step, we started by using the cluster tool of the GROMACS package with the gromos clustering algorithm [39] to obtain a representative (average) structure of each dimer. This initial step convoluted most of the conformational variability resulting from the MD relaxation step into 212 representative conformations.

To reduce even further the structural similarity among the conformations selected in the previous step, a second clustering procedure was applied. This second step required the calculation of an RMSD matrix (212*211/2) that accounts for the dissimilarity between dimer conformations. For the fit procedure and calculation of RMSD, we used the *β*-sheet core region to reduce the contribution of spurious structural variability into the RMSD. Finally, in the third step, we performed A/B to B/A monomer permutations to ensure that the best fits (lowest RMSD configurations) were chosen [40]. The final RMSD matrix was used with the clustering tool available in the HADDOCK 2.2 software [41, 42] to generate structurally homogeneous clusters at a RMSD cutoff value of 4 Å and using the gromos algorithm[39].

### 2.4. Calculation of binding energies with MM-PBSA

All binding free energy (*E_bind_*) calculations were performed using a new in-house implementation of the Molecular Mechanics Poisson-Boltzmann Surface Area (MM/PBSA) method [43, 44], written in the programming language Python (https://github.com/mms-fcul/mmpbsa). This method uses a single-trajectory approach to calculate 4 distinct binding energy components - 2 related to the Molecular Mechanics energy in vacuum, namely Van der Waals (*E_V dW_*) and Coulombic (*E_coul_*) terms, and 2 related to the Solvation energy, Polar (*Solv_polar_*) and Apolar (*Solv_apolar_*) - using the parameters of a specific force-field. In this method, *E_coul_* and *E_V dW_* are calculated using the equations representing the Coulomb and Lennard-Jones potentials, respectively. DelPhi4Py (https://github.com/mms-fcul/DelPhi4Py [29]), a DelPhi [45] python wrapper, is used as Poisson-Boltzmann (PB) solver to calculate the *Solv_polar_* energy. The final energy component, *Solv_apolar_*, is calculated using a widely used Solvent Accessible Surface Area Only model (SASA-Only) of the system. The *Ebind* is, thus, comprised of the sum of the 4 aforementioned energetic terms. The program is able to read simple GROMACS coordinate files (*.gro*), extracted from the MD final trajectories (*.xtc*), and calculate energies for each given frame.

In this work, GROMOS 54A7 compatible parameters (atomic charges, atomic radii [46] and 12-6 Lennard-Jones intermolecular pair potentials) were used for the calculation of the 4 energy components. In the calculations of *E_coul_* and *Solv_polar_*, a dielectric constant of 4 was used for the protein interior [47, 48]. In the Poisson-Boltzmann calculations, we used a grid scaling factor (*scale*) of 2 and a convergence criteria (*convergence*) of 0.01 kT/e [49]. We also used 500 and 50 linear (*nlit*) and non-linear (*nonit*) iterations, respectively. Additionally, the grid size (*gsize*) and the grid center (*acent*) were dynamically calculated for each file in the following manner: *gsize* was set as 2 times the maximum value of the atomic coordinates in the 3 axis, plus 1 if the final number is even; *acent* was set as the geometric center of all the atomic coordinates for a file. For each dimer, the MM-PBSA calculations were run on the full MD trajectories, every 100 ps, which resulted in 1000 frames per dimer.

## 3. Results

### 3.1. Stability analysis

An important goal of our work is to assess the stability of the 221 dimer conformations generated using the MC-ED protocol, and structurally characterise the most stable binding modes. However, since the MC-ED cost function creates a few atomic clashes and artificially introduces entanglements at the dimer interface, we designed a MM/MD-based protocol to relax the conformations before evaluating their energetic stability. In particular, we performed multiple steps of energy minimization followed by several initialization MD segments where position restraints were applied to each monomer with decreasing force constants. A set of 212 dimer configurations, were further relaxed using an unrestrained MD segment (100 ns), while the remaining 9 were discarded due to the presence of artificial entanglements (Figure S1 of Supporting Information). The goal of performing this long MD segment is to relax the dimer interfaces and potentially achieve better packing through local rearrangements.

For many dimers, a few nanoseconds was enough to relax their binding interfaces, while others, which were more dynamic, took a longer time to equilibrate. Several properties were used to monitor the equilibration procedure, including RMSD (vs. the initial and final structures), radius of gyration, secondary structure content, size of the interfacial area, etc (Figure S2 of Supporting Information). Since some dimers are more structurally unstable than others, we decided to take a conservative approach and discarded the initial 80 ns of the long MD segments. Therefore, all analyses performed and presented in what follows focused exclusively on the final 20 ns segments.

We observed significantly high values of the interfacial surface area in the ensemble of dimers comprising all relaxed conformations of the 212 MD simulations (Figure 2A), and an average value of ~8 *nm*^2^.

**Figure 2:**
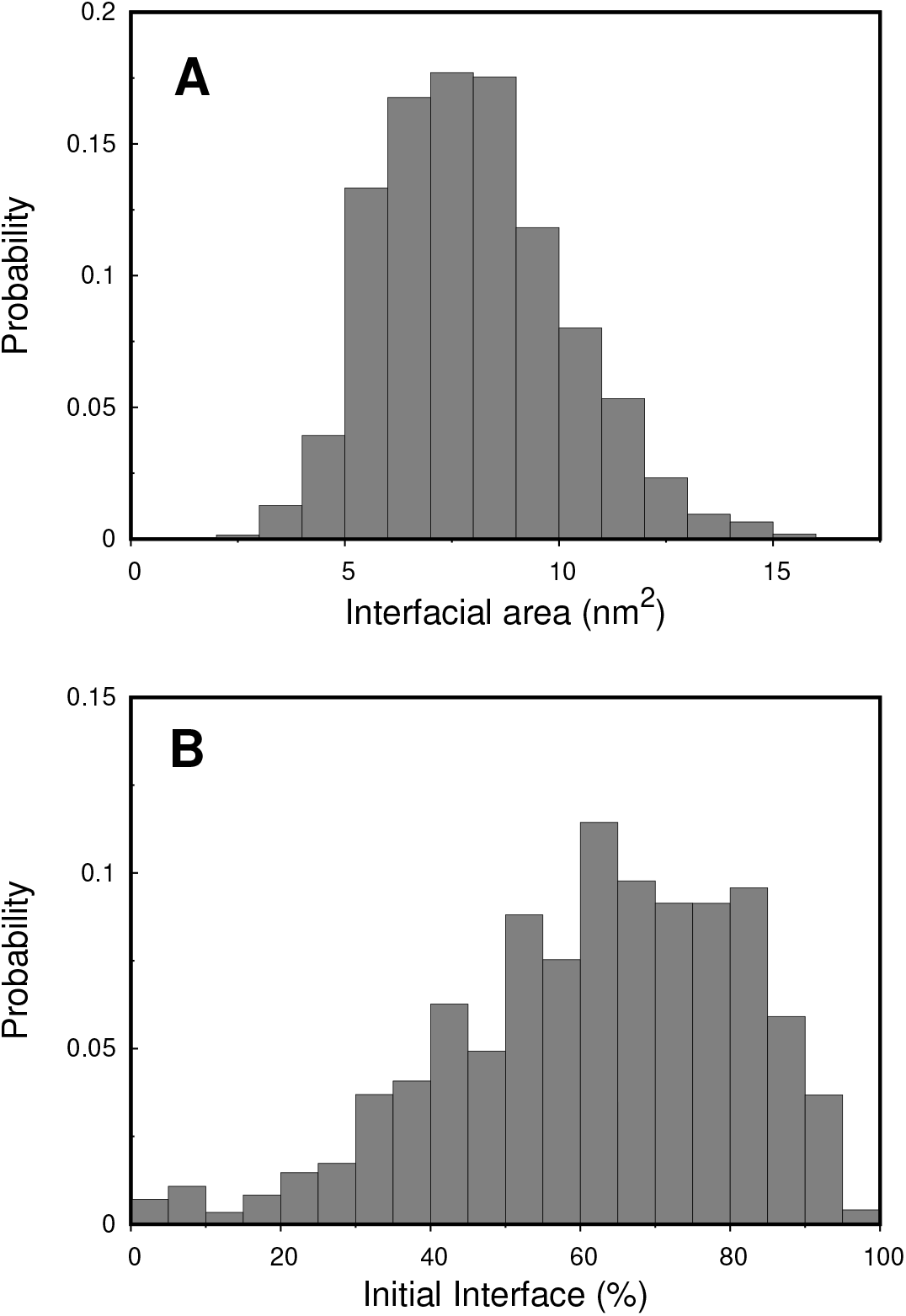
Interfacial area distribution (A) and percentage of initial interface (B) of the entire relaxed ensemble composed by the last 20 ns of the 212 MD simulations. The interfacial area shown is the average of the two area values mapped on each monomer. The residues of both monomers were included in the calculation of the initial interface percentage (see Methods for further details).

High interfacial area values are usually correlated with shape-complementarity, which is one of the major drivers of protein self-association taken into account in the docking protocol. Since all dimer conformations were relaxed during long MD segments, we also evaluated how much of their equilibrated interfaces diverged from the initial ones (Figure 2B). Most dimers (~ 75%) retained more than 50% of the original binding interface generated in the MC-ED, which suggests a high stability, and confirms the predicting ability of the MC-ED method.

To characterize the nature of the interactions governing/underlying dimer stability, we investigated the formation of cross-β structures or other less specific intermolecular hydrogen bonds (Figure S3 of Supporting Information). No significant differences were observed between the *β* structure content distributions of the full dimers and the sum of monomers, showing that the amount of cross-*β* is residual. The average number of intermolecular hydrogen bonds observed is also relatively modest with no preference between main chain and side chains.

Next, we evaluated dimer stability by means of the MM-PBSA approach, which provides binding energies. The binding energies calculated for the relaxed portion of our 212 MD simulations follow a normal distribution with an average ~-40 kcal mol^−1^ and a total range spanning from −100 to 0 kcal mol^−1^ (Figure 3A).

**Figure 3:**
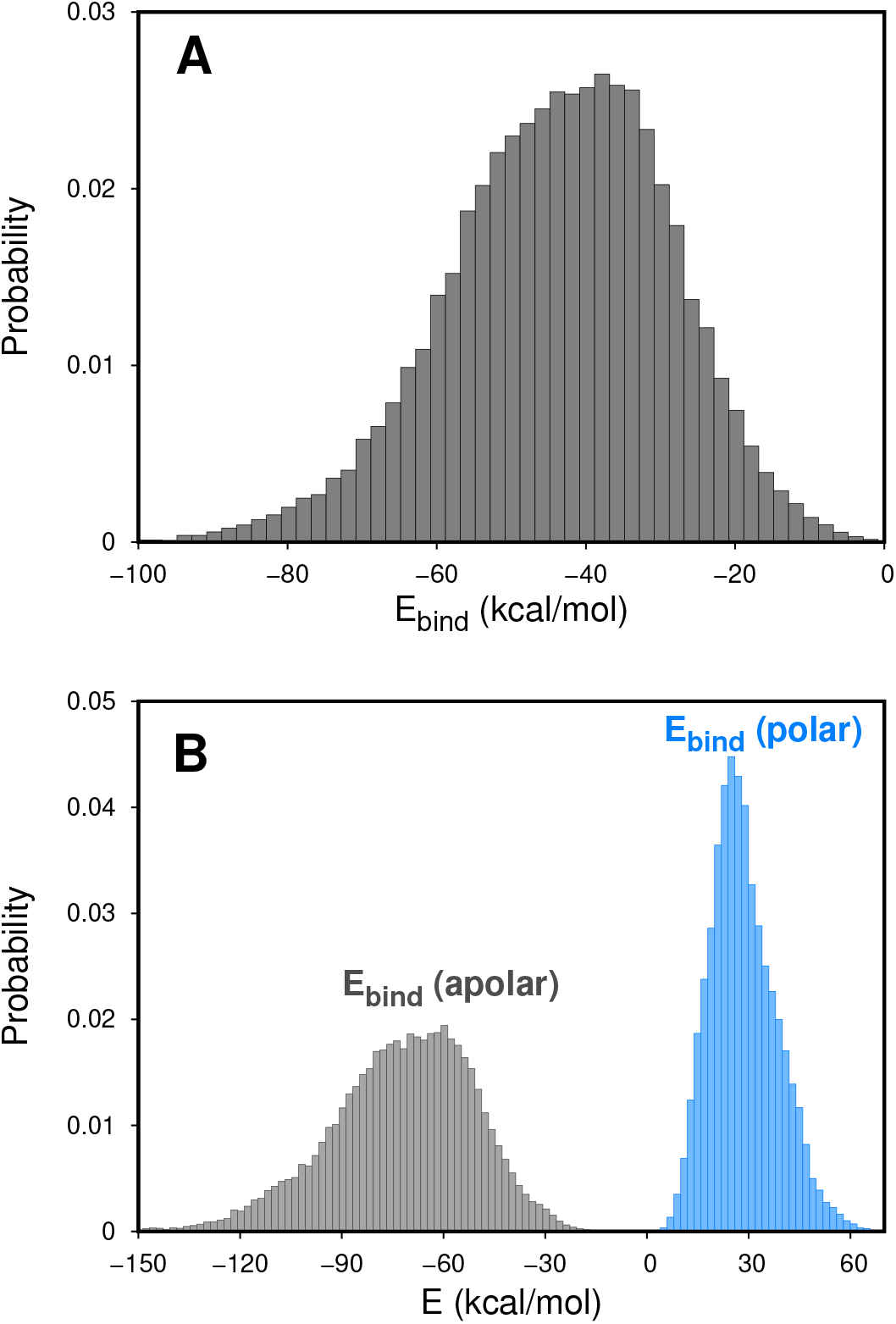
Probability histograms for the binding energies (A), and their polar (blue) and apolar (black) contributions (B) calculated with the MM-PBSA method over the complete ensemble of relaxed *β*_2_m dimers. The polar contribution of the binding energy is the sum of coulombic and polar solvation energies, while the apolar contribution was obtained from van der Waals and apolar solvation energies.

These low binding energies confirm that the interfaces obtained from MC-ED and relaxed using the MD protocol are very stable, which is consistent with the high interfacial area observed for most conformations. In the MM-PBSA formalism, the binding energy is usually decomposed in vacuum and solvation terms, or in polar and apolar contributions. By separating the polar from the apolar terms to the total energy, we observe that the apolar interactions are the main contributors to dimer stability (Figure 3B and Figure S4 of Supporting Information), suggesting that affinity and selectivity are not driven by polar contacts but rather by contact surface and shape complementarity.

We were intrigued by the positive polar energy contributions and tested the impact of changing the dielectric constant in the MM/PBSA methodology. This is an important parameter that despite being system-dependent, has been adjusted to fit different experimental data [48, 50, 51, 52] with the value of 4 leading to the best results [47, 48, 50, 51]. Nevertheless, we repeated the calculations of the polar energy contributions using *ϵ* = 3 and 5 (Figure S5 of Supporting Information). As expected from the Coulomb and Poison-Boltzmann formalisms, a higher dielectric constant leads to a larger attenuation between electrostatic interactions. Since these interactions are unfavorable, due to high energies, we observe better binding with *ϵ* = 5. However, while artificially increasing the dielectric constant can bring these contributions closer to zero, it will not make the interactions energetically favorable.

### 3.2. Structural analysis

#### 3.2.1. Identification of hot-spot residues

In protein aggregation the term ‘hot-spot” is often used to refer to residues (or regions) of the protein (typically non-polar groups that are buried inside the native structure) that give rise to intermolecular interactions between aggregation-prone intermediate states thus triggering the aggregation pathway. Here, we perform a fine-grained analysis of the dimer conformations in order to determine which residues are present at the dimer interface with higher probability, being potential candidates for aggregation hot-spots. The identification of hotspots resulting from this procedure differs from that adopted in previous studies [11, 12, 13] in which an hot-spot is a residue that establishes the most interactions within the ensemble of the most frequent interfacial contacts. An initial analysis was performed based on the relaxed conformations of all 212 dimers and the resulting probability histogram predicts several aggregation hot-spots that are located at both the N- and C-termini, as well as at the DE-loop (Figure S6 of Supporting Information). Nevertheless, this analysis does not allow one to distinguish between residues that actively stabilize the dimer interface from bystanders, i.e., residues that are highly likely to be present but do not establish any stabilizing role. With this idea in mind, we used the MM-PBSA data to sort all dimers in our ensemble according to their binding energies and constructed two data sets: one containing the best (i.e. more stable) interfaces, and another containing the worst (i.e. less stable) interfaces (10% cutoff). By subtracting from the probability of being present in the best (worst) interfaces *P_B_* (*P_W_*), the probability *P* mentioned above, we identified enriched regions of the protein (i.e. regions where the probability difference, Δ*P* =*P*_*B*(*W*)_ − *P* > 0) and impoverished regions of the protein (i.e. regions where the probability difference, Δ*P* =*P*_*B*(*W*)_ − *P* < 0) in the best and worst interfaces (Figure 4). For the most stable interfaces, the enriched regions comprise the N-terminus, the BC-loop, the DE-loop and the C-terminus. Interestingly, these regions correspond to the impoverished regions for the less stable interfaces, which further supports their stabilizing role in self-association. In contrast, for the less stable dimer interfaces the enriched protein regions comprise the CD-loop and EF-loop, which are the impoverished regions for the more stable interfaces. These results suggest that the more stable interfaces may trigger an aggregation pathway, which is different from that triggered by the less stable ones.

**Figure 4:**
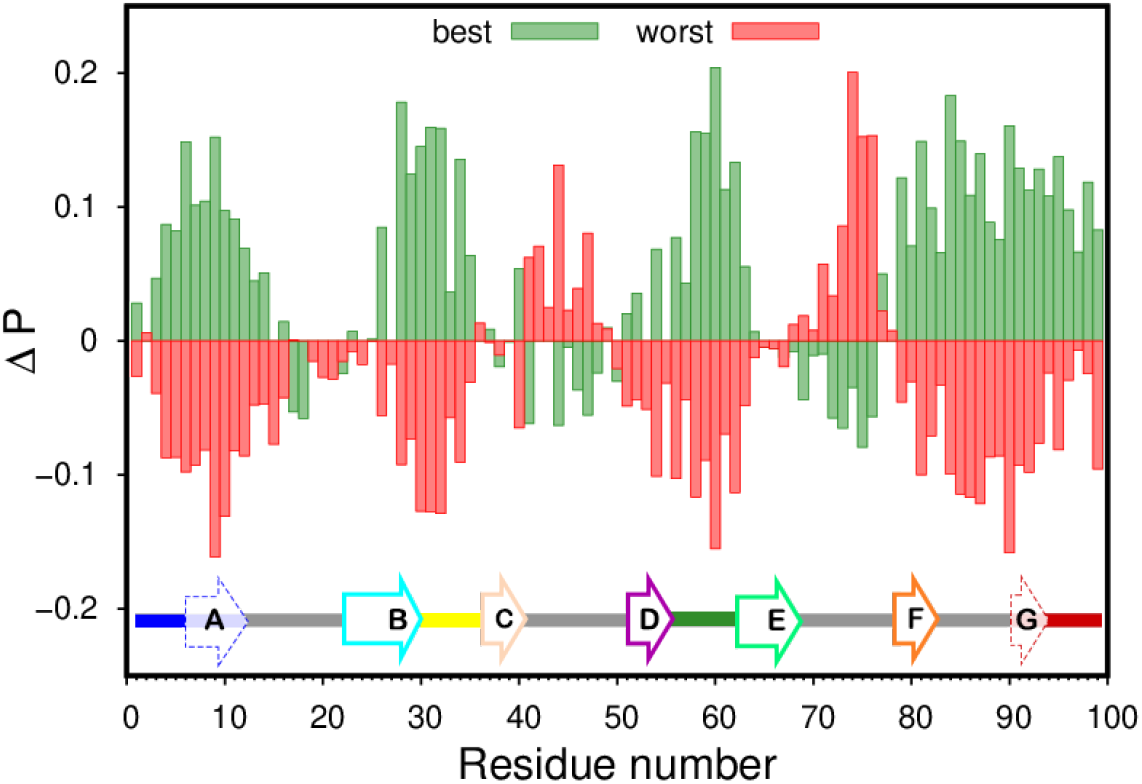
The most likely residues (positive probability difference; Δ*P*) in the interface of the most stable (green) and less stable (red) dimer interfaces. The best and worst dimer interfaces were selected based on a 10% cutoff on their binding energy. A linear depiction of the secondary structure content (bottom) using the color code of Figure 1. *β* sheets A and G have a dashed arrow and are included into their adjacent termini since these domains are significantly unstructured in the I2 *β*_2_m intermediate [13].

#### 3.2.2. Identification and structural characterization of stable binding modes

After determining potential aggregation-prone regions of the D76N mutant of *β*_2_m, we set out to identify the most stable binding modes (BMs). We started by applying a clustering protocol to ensure that only unique interfaces, corresponding to different BMs were selected. By using this approach, we obtained 161 individual BMs (clusters) from the 212 representative dimer conformations. Using the MM-PBSA binding energy values, we ranked these BMs according to their stability, and the top scorers were used in further analyses (Table 1).

**Table 1:** Binding energies and dimer interface details for the top 10 best clusters/binding modes. The binding energy contributions were obtained directly from the MM-PBSA calculation and the interfacial area was calculated using our SASA-based protocol. The initial interface percentage was obtained comparing the presence of the individual residues at the dimer interface after the MD relaxation step in relation to the initial structure.

| Binding Mode | $E_{bind}$<br>(*kcal/mol*) | $E_{bind}$ (Apo)<br>(*kcal/mol*) | $E_{bind}$ (Pol)<br>(*kcal/mol*) | Interf.<br>Area (*nm*<sup>2</sup>) | Initial<br>Interf. (%) |
| --- | --- | --- | --- | --- | --- |
| 1 | -82.9 ± 7.1 | -137.0 ± 10.3 | 54.1 ± 4.5 | 14.2 | 73.4 |
| 2 | -82.1 ± 5.2 | -121.5 ± 6.5 | 39.4 ± 2.5 | 12.9 | 60.6 |
| 3 | -79.0 ± 3.2 | -115.4 ± 2.8 | 36.5 ± 1.5 | 11.4 | 92.1 |
| 4 | -77.3 ± 7.6 | -104.7 ± 12.5 | 27.4 ± 5.0 | 10.7 | 5.8 |
| 5 | -75.4 ± 1.3 | -106.0 ± 1.0 | 30.6 ± 0.3 | 10.9 | 78.4 |
| 6 | -71.8 ± 13.4 | -121.9 ± 18.8 | 50.1 ± 5.9 | 13.2 | 82.9 |
| 7 | -71.0 ± 6.9 | -122.8 ± 10.3 | 51.8 ± 3.7 | 13.7 | 62.5 |
| 8 | -68.0 ± 3.3 | -110.3 ± 6.9 | 42.3 ± 3.9 | 11.4 | 83.6 |
| 9 | -67.5 ± 3.2 | -116.5 ± 5.1 | 49.0 ± 3.0 | 12.7 | 66.5 |
| 10 | -67.2 ± 2.0 | -105.9 ± 1.1 | 38.7 ± 1.0 | 10.1 | 66.2 |

In the 10 most stable binding modes captured by our protocol, we noticed BM-4 as an outlier since the percentage of initial interface upon relaxation was overwhelmingly low, as a result of a large conformational rearrangement. Even though this BM was not predicted by the MC-ED, the high stability still makes it worth investigating. Overall, the top BMs obtained from our clustering analysis have low binding energies spanning from −83 to −67 *kcal/mol* and, as expected, they also display high interfacial areas, with the most stable BM having the highest interfacial area. However, there is no clear correlation between the size of the interface and the final binding energies, indicating that the nature of the residues involved in the intermolecular interactions is also important. We used these 10 BMs to calculate a probability histogram for the presence of residues in the interfacial region, which confirmed the results reported previously that considered the ensemble of 212 dimers (Figure S7 of Supporting Information).

The three-dimensional structural representation of the most stable BMs highlights a remarkable diversity of dimerization interfaces (Figure 5), which suggests a complex and heterogeneous oligomerization pathway for the D76N mutant of *β*_2_m. By visual inspection, one can easily recognize the aggregation-prone regions (highlighted as colored patches) that were previously identified (Figure S7 of Supporting Information). In particular, the N- or the C-ter are present in the interfaces of all the binding modes reported in Figure 5. Since the BC/DE loops are usually located on the same side of the monomeric form of *β*_2_m, binding through this interface (as in the case of BM-1, 6, and 7) should not be limited two these structural elements to allow for further oligomerization. Therefore, it is likely that the two terminal regions will play a role in the oligomerization process, by providing alternative binding modes.

**Figure 5:**
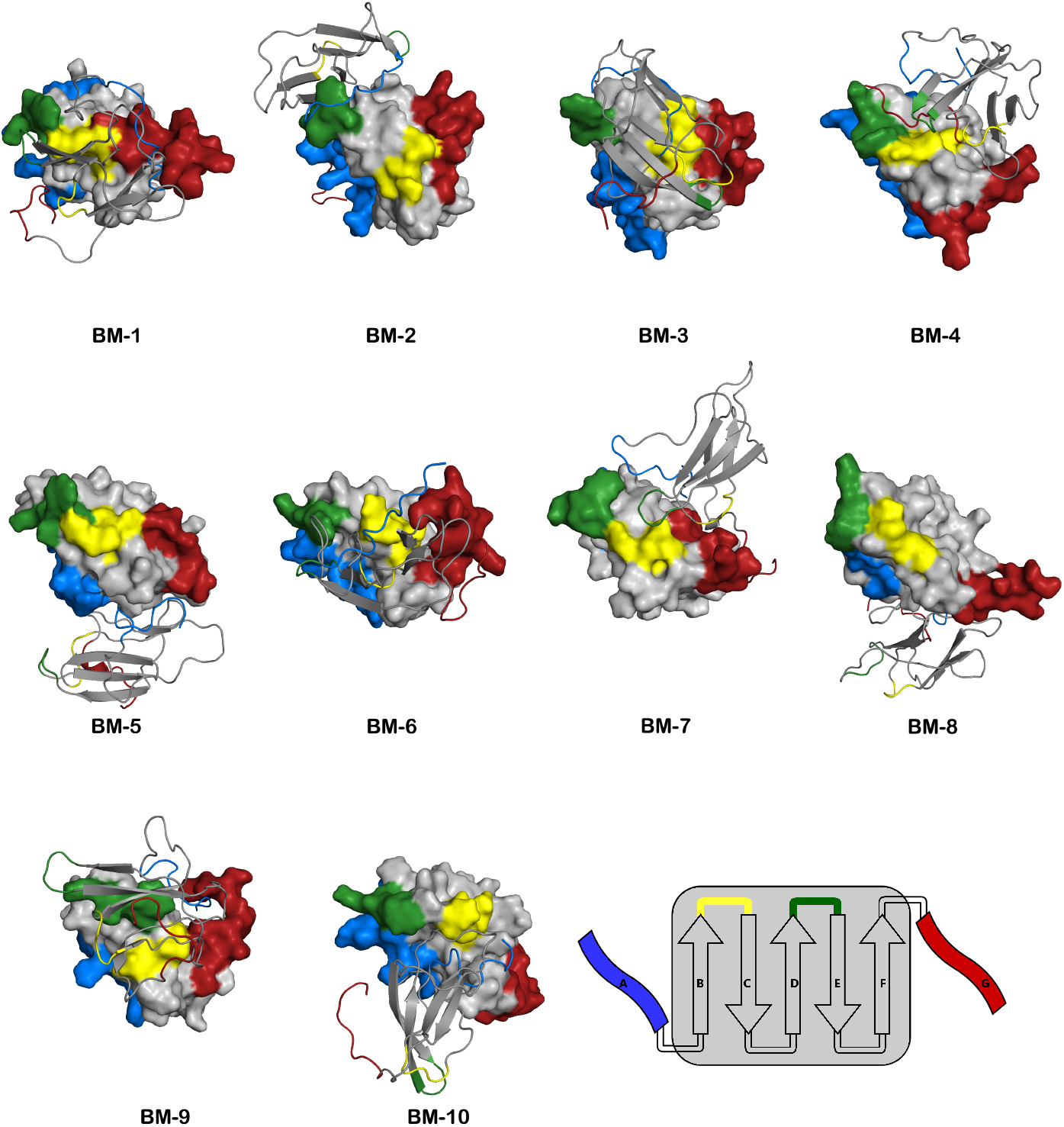
Structural representation of the 10 most stable *β_2_*m dimer binding modes. A schematic representation of the secondary structure and color code is also shown. The N- and C-terminus, and the BC- and DE-loop are colored in blue, red, yellow and green, respectively. The structures are aligned by the monomer represented with a surface, while the other monomer is represented with cartoon.

#### 3.2.3. Oligomerization of stable binding modes

In order to explore how early protein–protein interactions may determine the formation of specific aggregate arrangements, we designed a simple molecular visualization protocol that generates larger oligomers by propagating the initial dimerization interface of a selected BM. In particular, we generated 6-mers by adding 4 dimeric units to a starting dimer representative of a stable BM (Figure 6).

**Figure 6:**
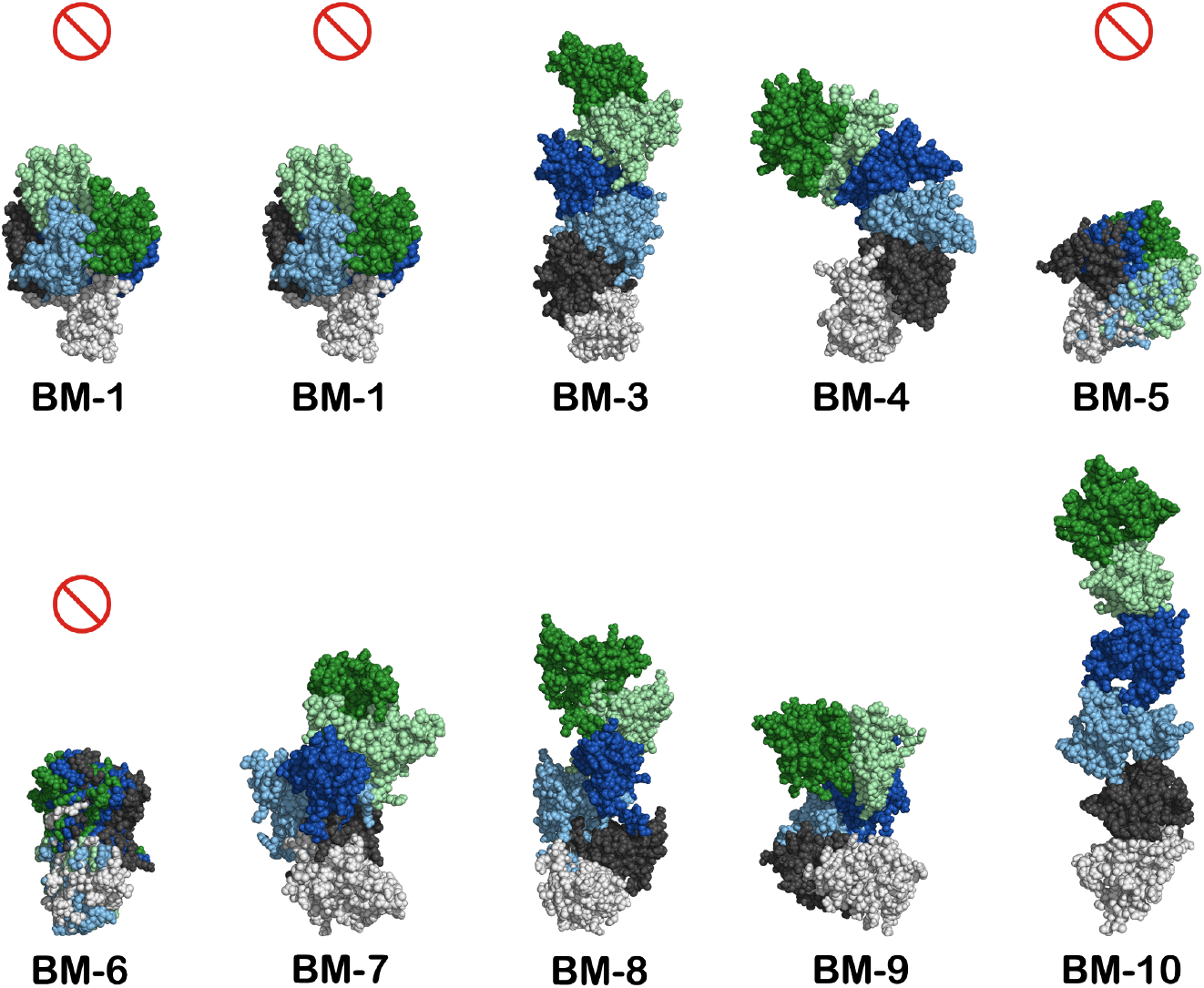
Oligomerization process for the 10 best BMs. Three dimensional representation of 6-mer oligomers where the interface of the initial dimer (MonA/B in white/black spheres, respectively) was propagated by adding four new dimer units via the alignment of the new MonA to the previous MonB (using Pymol[38]). With the exception of the first dimer where MonA is colored in light grey, all MonA conformations were hidden, which results in a 6-mer with the ABBBBB monomer arrangement. The progression of the oligomerization (MonBs) follows the color code: black; light blue; blue; light green; green. A prohibition sign indicates the limited growth BMs.For visual clarity we only report 6-mers but the structure of larger oligomers can be easily inferred.

The oligomers thus obtained can be classified into limited and unlimited growth oligomers, depending on whether the addition of more units leads to an oligomeric chain that closes upon itself, or, in alternative, a chain that may sustain further grow. Limited growth oligomers are usually obtained when the interfaces of each monomer are oriented in a such a way that the addition of further units leads to closed structures. The simplest case is a head-to-head arrangement, in which both monomers have the same residues in the interface. BMs 5 and 6 are examples of limited growth dimers in which both monomers interact via the same regions (BM-5: Nter+CDloop+Cter; BM-6: BCloop+DEloop+Cter). BM-1, the most stable binding mode reported here, leads to closed trimers with an interface involving the Nter+BCloop+DEloop from one monomer and the BCloop+DEloop+Cter from the other.

The unlimited growth oligomers can be further classified into linear (BMs 2, 3, 8, and 10) and helical oligomers (BMs 4, 7, and 9). All these BMs have contact interfaces where the monomers interact via different regions and residues. This leads to the generation of free contact surfaces that permit further oligomerization. The relative orientation of monomers at the interfaces is what determines the formation of linear or helical aggregates. For these binding modes, only a small rearrangement of the interfacial/contact regions is required to accommodate another monomer, which suggests an efficient oligomerization process.

This simple analysis shows that, in the absence of major conformational rearrangements, the relative orientation of the monomers in the initial interfacial region is thus key to determine the shape and size of the final aggregates. The limitation to grow beyond a certain size can be overcome by mixing different BMs, or if the final oligomer is prone to undergo a major conformational change compatible with further growth.

### 4. Discussion and Conclusion

Protein aggregation is a remarkably complex process, where the formation of molecular structures of varying size and timescales is highly dependent on environmental conditions. Solving the aggregation mechanism, requires determining the size, distribution and structures of the oligomeric assemblies, filaments, protofibrils and fibrils that populate the amyloid pathway, as well as the rate constants associated with every transition [53]. In the case of *β*_2_m it is even a bigger challenge since the wild-type form does not aggregate *in vitro* under physiological conditions. Despite these difficulties, progress has been made regarding the initial phase of the aggregation mechanism (reviewed in [10]), as well as in determining the structure of amyloid fibrils [54]. In particular, research carried out by several research groups indicate that the first phase of the *β*_2_m aggregation mechanism is the dimerization of aggregation prone monomers [10].

Here, we studied in detail an ensemble of dimers that resulted from the association of an aggregation-prone monomeric state populated by the D76N natural mutant of *β*_2_m. This mutant has attracted considerable attention in the last years because, contrary to the WT form, it aggregates readily *in vitro* in physiological conditions [8, 21]. The monomeric intermediate state was previously reported in a molecular simulation study based on discrete molecular dynamics simulations of a full-atomistic structure-based model [12, 13, 55]. It is structurally characterized for having the two terminal regions unstructured and detached from a well-preserved core. An intermediate state sharing the same topology was later reported in a study that combined ss-NMR with Molecular Dynamics simulations [20].

The ensemble of dimers investigated here was prepared with an in house protein-protein docking protocol that optimizes the interfacial region for shape, electrostatic and hydropathic complementarity [13]. While the docking method provides a qualitative characterization of the dimerization interfaces, it cannot be used to assess their stability in a quantitative manner. For this reason, in the present study we designed a simulation protocol that combines classical Molecular Dynamics with MM-PBSA to relax the dimer interfaces and accurately estimate their binding energies. We further designed a structural clustering protocol that allowed us to isolate unique dimer configurations representative of distinct binding modes. Our analysis shows that the major driving force of interface stabilization are interactions between apolar residues. This is perhaps not surprising taking into account the fact that in its functional form *β*_2_m is docked onto the *β*3 domain of the major histocompatibility complex (MHC-I) through the four-stranded (ABED) beta-sheet, and that the four aromatic, bulky residues Phe56, Trp60, Phe62, and Tyr63 and the aliphatic Leu65, which are shielded from the solvent in the quaternary structure of the MHC, become highly solvent-exposed in the monomeric form, thus being able to act as sticky patches in intermolecular association. In line with this observation, we find that the interfaces of the most stable binding modes are particularly rich in residues pertaining to both the N- and C-terminus, as well residues from the BC- and DE-loops. We highlight Trp60 from the DE-loop, which is classically recognized as an aggregation hot-spot for *β*_2_m[10]. On the other hand, the less stable interfaces are stabilized by intermolecular interactions involving residues from the CD- and EF-loops.

Since there is no clear relation between dimer stability and aggregation potential [56], in the present work, instead of focusing on the most stable dimer, we considered an ensemble of dimers. The three-dimensional structural representation of the most stable binding modes revealed a remarkable variability of dimerization interfaces. In order to investigate the early protein–protein interactions that establish at the dimer level and can determine the specific aggregate arrangements, we devised a simple geometric rule combined with a visualization protocol that propagates the interfaces of the 10 most stable binding modes. Perhaps not surprisingly, we found that in the absence of major conformational rearrangements, the relative orientation of the monomers in the initial interfacial region is key to determine the shape and size of the final aggregates. Moreover, the dimerization interfaces in which the participating monomers contribute with the same residues (e.g. in a head-to-head arrangement) limit the growth of relatively small oligomers, while more heterogeneous dimeric interfaces in which monomers interact via different regions and residues have a potential to be propagated leading to larger oligomeric chains.

## Conflicts of interest

There are no conflicts of to declare.

## Author contributions

MM and PFNF designed research; NFBO and FEPR performed calculations; JNMV developed software; RJSL generated the initial data for MD simulations; NFBO and FEPR prepared the figures; all authors analyzed the data; NFBO, FEPR and JNMV prepared a preliminary version of the manuscript; PFNF and MM wrote final version.

## Acknowledgements

We acknowledge Diogo Vila Viçosa, Pedro Reis, and Tomás Silva for valuable discussions. António M. Baptista is also acknowledged for developing the fixbox tool, which was used to correct the periodic boundary conditions in the dimer trajectories. We acknowledge financial support from Fundação para a Ciência e a Tecnologia through grant CEECIND/02300/2017 and projects PTDC/FIS-OUT/28210/2017, UIDB/04046/2020, and UIDP/04046/2020.

## Notes

### Competing Interest Statement

The authors have declared no competing interest.

