## Supplemental information for "Predicting stable binding modes from simulated dimers of the D76N mutant of *β*2-microglobulin"

### Supporting Information

July 14, 2021

### List of Figures

Figure S1: Cartoon representation of a structurally linked dimer. . . . . . . . . . . . . . . . . . S3

Figure S2: Equilibrium properties from the MD simulations . . . . . . . . . . . . . . . . . . . . . . . S4

Figure S3: $\beta$-structure content and H-bonds distributions. . . . . . . . . . . . . . . . . . . . . . . . . S5

Figure S4: Binding energy contributions scatter plots . . . . . . . . . . . . . . . . . . . . . . . . . . . . S5

Figure S5: Energy distributions of the polar and apolar contributions. . . . . . . . . . . . . S6

Figure S6: Probability histogram of interface residues in the complete ensemble. . . S7

Figure S7: Probability histogram of interface residues in the top BMs. . . . . . . . . . . . . S8

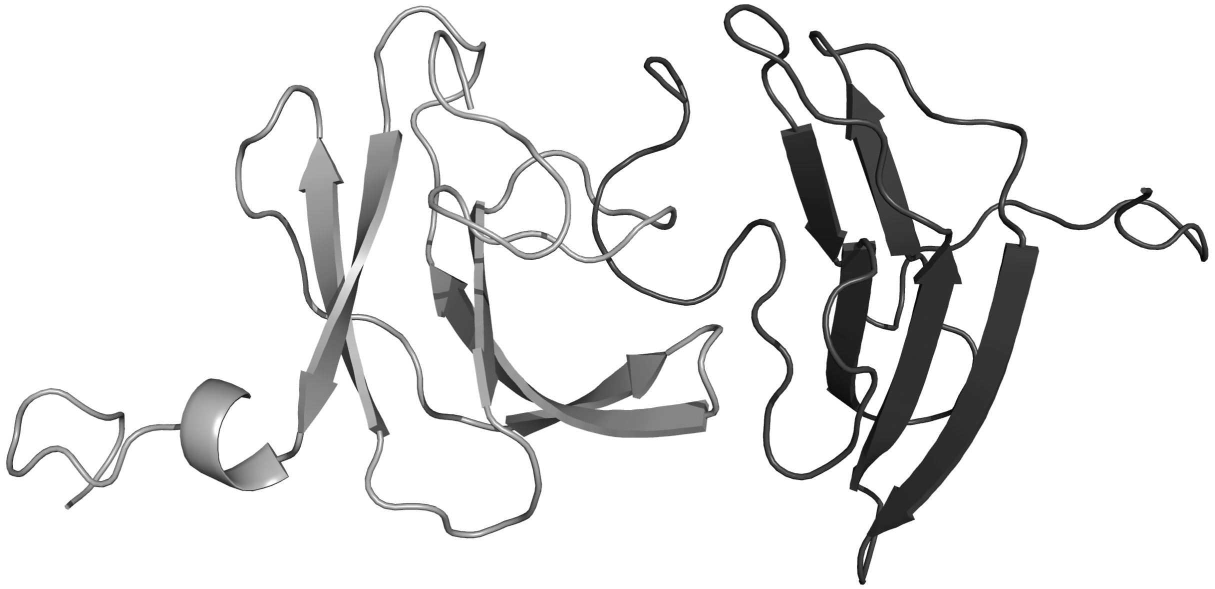

Figure S1: Cartoon representation of a structurally linked dimer with one monomer coloured in grey and the other in dark grey. From the initial 221 dimers, 9 displayed this type of partially entanglement introduced by the MC-ED protocol, and were discarded.

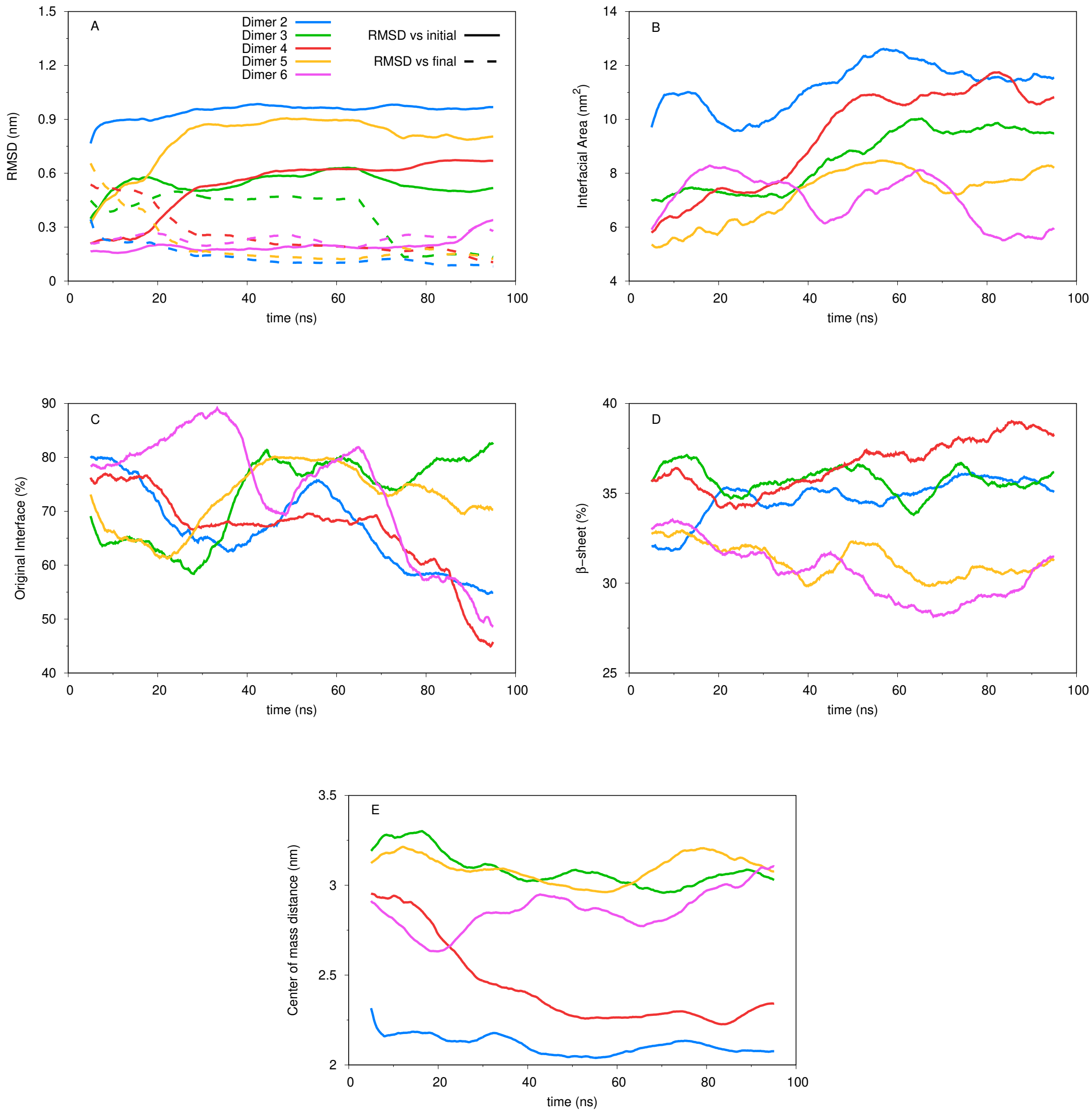


Figure S2: RMSD values compared to the initial and final conformations of the dimers (A), interfacial area (B), percentage of initial interface (C), $\beta$-sheet content (D), and distance between the centers of mass of both monomers (E) for the first 5 dimers. Taking into consideration the significant size of our full ensemble (212 dimers in total), we show in detail only a small representative sample of the data we have analyzed.

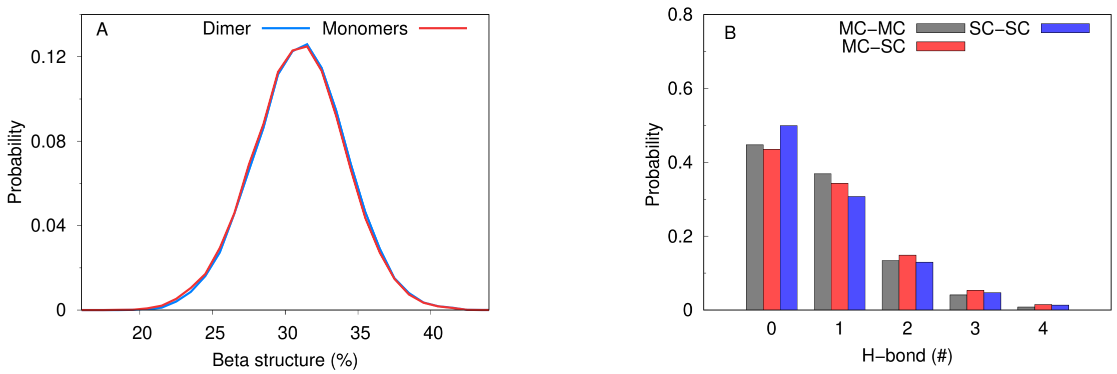


Figure S3: Probability distributions of $\beta$ structure content ($\beta$-sheet and $\beta$-bridge from the DSSP criteria) obtained for both monomers and dimer (A), and of the number of intermolecular hydrogen bonds, splitted between main-chains and side-chains (B). All data used was obtained from the equilibrated/relaxed ensembles, corresponding to the last 20 ns of each of the 212 dimer simulations.

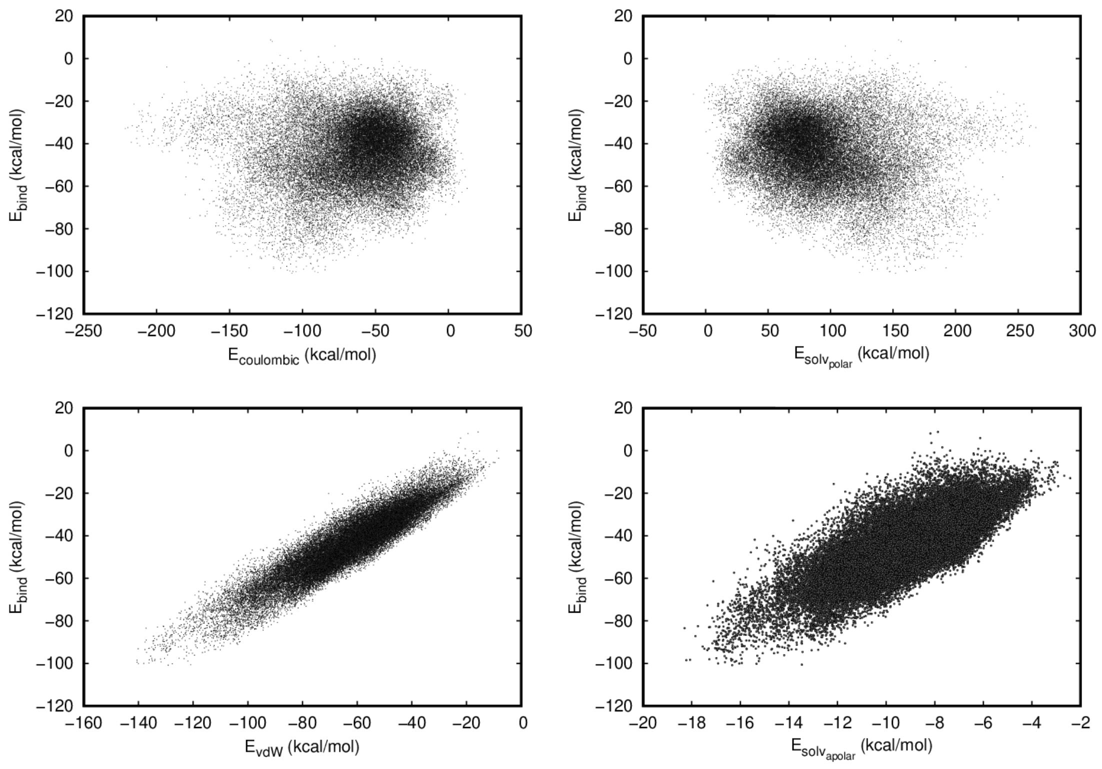


Figure S4: Scatter plots of the binding energy correlated with coulombic (A), polar solvation (B), van der Waals (C), and apolar solvation (D) contributions. These energy values were obtained from the relaxed ensembles using the MM-PBSA method.

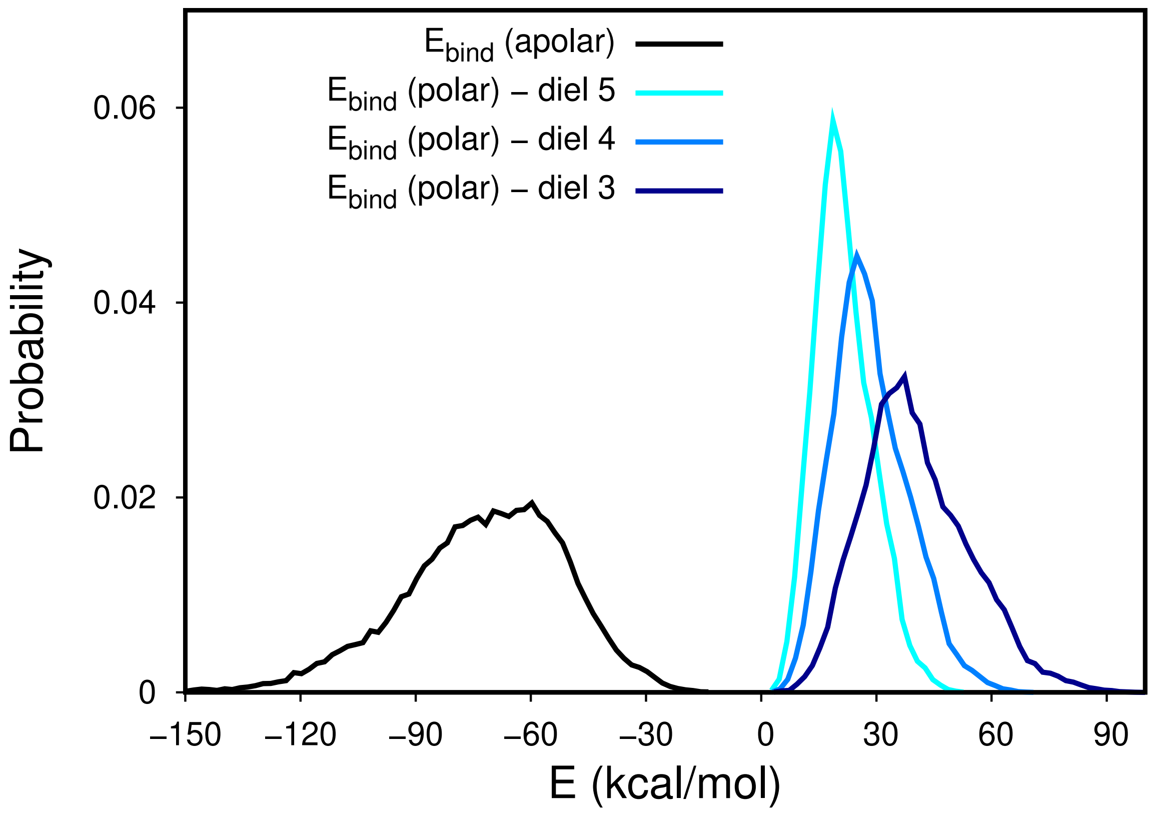


Figure S5: Energy distributions of the polar and apolar contributions calculated from the relaxed ensembles using the MM-PBSA method. The polar contribution was calculated using three different dielectric constant values, namely 3, 4 and 5, for the protein interior.

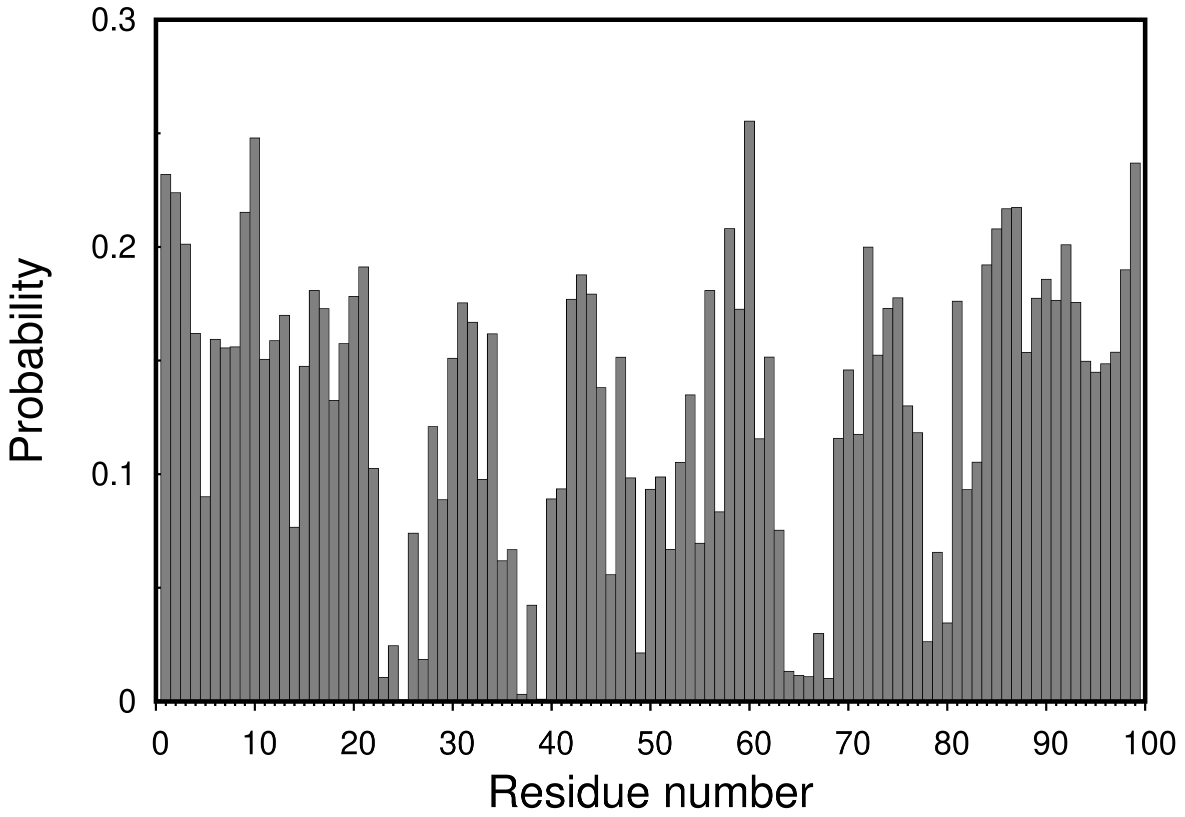


Figure S6: Probability histogram of the interfacial residues from the complete relaxed ensemble of dimer configurations. The probability of each residue being at the interface was calculated from their interfacial area values and applying a cutoff (see Methods).

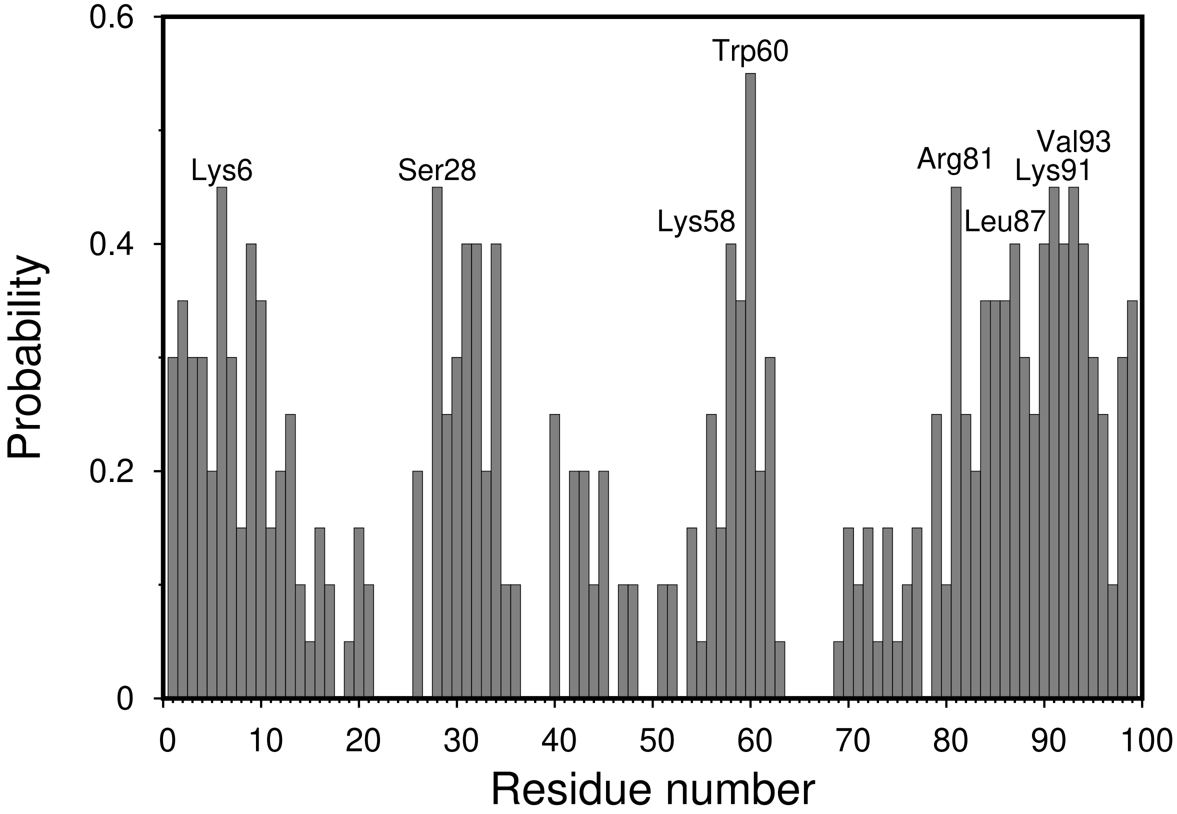


Figure S7: Probability histogram of the interfacial residues in the 10-best binding modes. The probability of each residue being at the interface was calculated from their interfacial area values and applying a cutoff (see Methods).Residues with higher probability of being in the interface have been shown above their respective histogram bars.
